# Different salicylic and jasmonic acids imbalances are involved in the oxidative stress-mediated cell death, induced by fumonisin B1 in maize seedlings with contrasting resistance to *Fusarium verticillioides* ear rot in the field

**DOI:** 10.1101/2019.12.25.882597

**Authors:** Santiago N. Otaiza-González, Verónica S. Mary, Silvina L. Arias, Lidwina Bertrand, Pilar A. Velez, María G. Rodriguez, Héctor R. Rubinstein, Martín G. Theumer

**Author notes:** Santiago N. Otaiza-González; Verónica S. Mary; Silvina L. Arias; Lidwina Bertrand; Pilar. A. Velez; María G. Rodriguez; Héctor R. Rubinstein; Martín G. Theumer. Corresponding Author Martín G. Theumer, Ph.D. Centro de Investigaciones en Bioquímica Clínica e Inmunología (CIBICI), UNC, CONICET, Departamento de Bioquímica Clínica, Facultad de Ciencias Químicas, Universidad Nacional de Córdoba, Haya de la Torre y Medina Allende, Ciudad Universitaria, X5000HUA, Córdoba, Argentina. Phone: (+54) (Ext. 3146). Fax: (+54).

## Abstract

**Background and aim:** Fungal and plant secondary metabolites modulate the plant-pathogen interactions. However, the participation of fumonisins in the *Fusarium verticillioides*-maize pathosystem is unclear. In this work was studied the cell death, and the reactive oxygen species (ROS) - phytohormone imbalance interplay underlying the phytotoxicity of fumonisin B1 (FB1) in maize germplasms with contrasting resistance to *Fusarium* ear rot in the field.

**Methods:** Resistant (RH) and susceptible hybrid (SH) maize seedlings, grown from uninoculated seeds irrigated with FB1 (1 and 20 ppm), were harvested at 7, 14 and 21 days after planting, and were examined for electrolyte leakage (aerial parts); and for oxidative stress biomarkers (aerial parts and roots). The phytohormone (salicylic and jasmonic acids) imbalance interplay underlying the FB1-induced cell death were further explored in seedlings exposed 24 h to the mycotoxin (1 ppm) in hydroponics.

**Results:** Cell death increased in RH and SH watered with 1 and 20 ppm of mycotoxin, respectively. Both toxin concentrations were pro-oxidant, and the major perturbations were found in roots. An Integrated Biomarker Response index was calculated suggesting that phytotoxicity occurs in a redox context more efficiently controlled by RH.

**Conclusion:** The pre-treatment with the antioxidant ascorbic acid led to the conclusion that cell death in RH was related to a salicylic acid increase mediated by ROS. Nevertheless, FB1 induced two different phytohormonal regulatory mechanisms mediated by oxidative stress in both maize hybrids.

## Introduction

Fumonisins are a set of mycotoxins primarily produced by the secondary metabolism of toxicogenic strains of *Fusarium*, mainly *F. verticillioides* and *F. proliferatum*; even though, in recent years, it has also been observed that these fumonisins can be synthesised by some black aspergilli (Frisvad *et al*., 2011; Susca *et al*., 2014). The group includes fumonisin analogs belonging to four primary series (FA, FB, FC and FP), although fumonisin B1 (FB1) is undoubtedly the most relevant because of its incidence in maize and its toxicity to human beings and animals.

*F. verticillioides*, an hemibiotrophic fungus that requires living plant cells in its early stages of colonization, infects maize all over the world causing severe pathologies such as ear, stem, root and grain rot. This fungus attacks stalks, kernels, and seedlings in all stages of development, inducing pre- and post-harvest diseases. Sometimes the damage remains unnoticed, and the infection can spread to the root system and cause seedling underdevelopment. Under certain conditions, it causes root and stem rot, increasing the possibility of overturning. Diseases are the result of complex interplays of environmental conditions, and the intrinsic characteristics of both the pathogen and the host (CAST, 2003). About the latter, maize germplasms generally respond differently to the infection by *Fusarium* spp.; some of them are susceptible, while others exhibit greater resistance to the fungal phytopathology (Santiago *et al*., 2015).

A large number of low-molecular-weight secondary metabolites synthesised by both the fungi and the plants may be involved, at some extent, in the outcome of the plant-pathogen interactions (Pusztahelyi *et al*., 2015; Selin *et al*., 2016). While some fungal metabolites are essential for virulence over specific plants, others act as non-host selective toxins that may contribute to pathogenicity. The F*. verticillioides*-maize link at the molecular level is not known in depth; however, several plant and fungal substances, including FB1, must be involved in the biochemical communication in both senses. The toxicodynamics of this mycotoxin seems to be, at least, partially shared in animals and plants, and it would be mainly related to the competitive inhibition of the toxin over the ceramide synthase activity, leading to imbalances in cellular lipids that have structural functions, and are involved in cell signalling (IPCS-WHO, 2000). In a previous work, we found that FB1 induced contrasting lipid imbalances depending on the hybrid resistance-susceptibility to the *F. verticillioides* invasion, mimicking those found in the fungal infection. The toxin significantly raised the spinganine (Sa) and the phytosphingosine (Pso) levels in resistant (LT 622 MG, RH) and susceptible (HX 31P77, SH) maize hybrids. However, in RH, the FB1 induced a greater increase of Sa, whereas in SH, higher levels of phytosphingosin (Pso) were observed, and it was speculated that the Sa increase would favour the pathogen elimination by activating localised cell death pathways (Arias *et al*., 2016).

Maschietto and collaborators showed the induction of oxidative stress in ears of resistant and susceptible maize lines inoculated with *F. verticillioides* (Maschietto *et al*., 2016), and FB1 is probably involved in such outcome. The oxidative stress was induced as a plausible mechanism for the FB1 toxicity in animal and plant cells (Wang *et al*., 2016; Xing *et al*., 2013). Studies performed in *Arabidopsis thaliana* pointed out the involvement of reactive oxygen species (ROS) as chemical mediators of lipid-induced cell death, whose levels are increased by exposure to FB1 (Saucedo-Garcia *et al*., 2011). Moreover, Zhao and co-workers (2015) showed that ROS accumulation caused by FB1 was reduced by breakdown products of indole glucosinolate with antioxidant behaviour.

Despite the fact that several studies showed the phytotoxicity of FB1, the data available about the involvement of this mycotoxin in the phytopathogenesis of maize diseases by *F. verticillioides* are not conclusive. For instance, there is differing information regarding the distribution of the toxin in plants. While some studies suggested that the fungus-plant interaction is necessary for FB1 translocation in maize seedlings (Zitomer *et al*., 2010), in a recent work conducted by our group, it was observed that the toxin disseminated to the aerial parts of the maize plants when administered via watering (Arias *et al*., 2016).

Regardless of the toxin distribution throughout the plants, symptoms indicative of disease induced by *F. verticillioides* were found in maize seedlings grown from uninoculated seeds irrigated with FB1 solutions (Arias *et al*., 2012; Williams *et al*., 2007), showing that the toxin is probably involved in the pathogenicity of this fungal infection. Moreover, Glenn and collaborators (2008) reported that the ability to develop foliar disease symptoms on maize seedlings by FB1 non-producing strains of *F. verticillioides* was restored in fumonisin-producing transformants, therefore indicating that the toxins contribute to the fungal pathogenesis. Conversely, other studies suggest that *F. verticillioides* do not require the synthesis of fumonisins to cause maize root and ear infections, or to produce ear rot (Dastjerdi and Karlovsky, 2015; Desjardins and Plattner, 2000). Therefore, further research must be conducted in order to elucidate the participation of fumonisins in the *F. verticillioides* invasion and pathogenesis in maize as well as the mechanisms underlying their effects.

Previous studies show that FB1 is an inducer of cell death (Asai *et al*., 2000; Igarashi *et al*., 2013; Glenz *et al*., 2019) by mechanisms not fully elucidated. In this regard, salicylic acid (SA) is a phytohormone commonly associated with the positive regulation of hypersensitive response-type cell death. It has a central role in defence and induces the activation of pathogenesis-related genes (PR), which generates resistance to a wide range of pathogens (Klessig *et al*., 2018; Loake and Grant, 2007). The cell death induced by FB1 in *Arabidopsis* is dependent on both, the accumulation of ROS and the synthesis of SA (Xing, 2013). Jasmonic acid (JA) may contain the spread of lesions caused by ROS, having this phytohormone an antagonistic effect on SA (Overmyer *et al*., 2003). However, Zhang *et al*. (2015) showed that the signaling pathway of JA is inhibited by FB1. Despite these studies show a central role of SA in the phytotoxicity of FB1 in *Arabidopsis*, the cell death induced by this mycotoxin, and how ROS and phytohormones, such as SA and JA, modulate this process, must still be explored in depth in plants of agronomic interest such as maize.

In this work was studied the cell death, and the reactive oxygen species (ROS) - phytohormone imbalance interplay underlying the phytotoxicity of FB1 in seedlings of maize hybrids with contrasting resistance to *Fusarium* ear rot in the field.

## Results

### Phytotoxicity in maize seedlings watered with FB1 Conductivity

Different profiles of cell death were observed between hybrids and levels of exposure to FB1 (1 and 20 ppm). The electrolyte leakage decreased at 14 dap in SH watered with 1 ppm of FB1, and increased in RH, at the same endpoint and mycotoxin concentration (Fig. 1). These alterations were transient, since conductivities remained unaltered in both hybrids at 21 dap. The highest toxin level tested (20 ppm) increased cell death in SH at 21 dap, but had no previous effects on this hybrid or on RH.

**Fig. 1.**
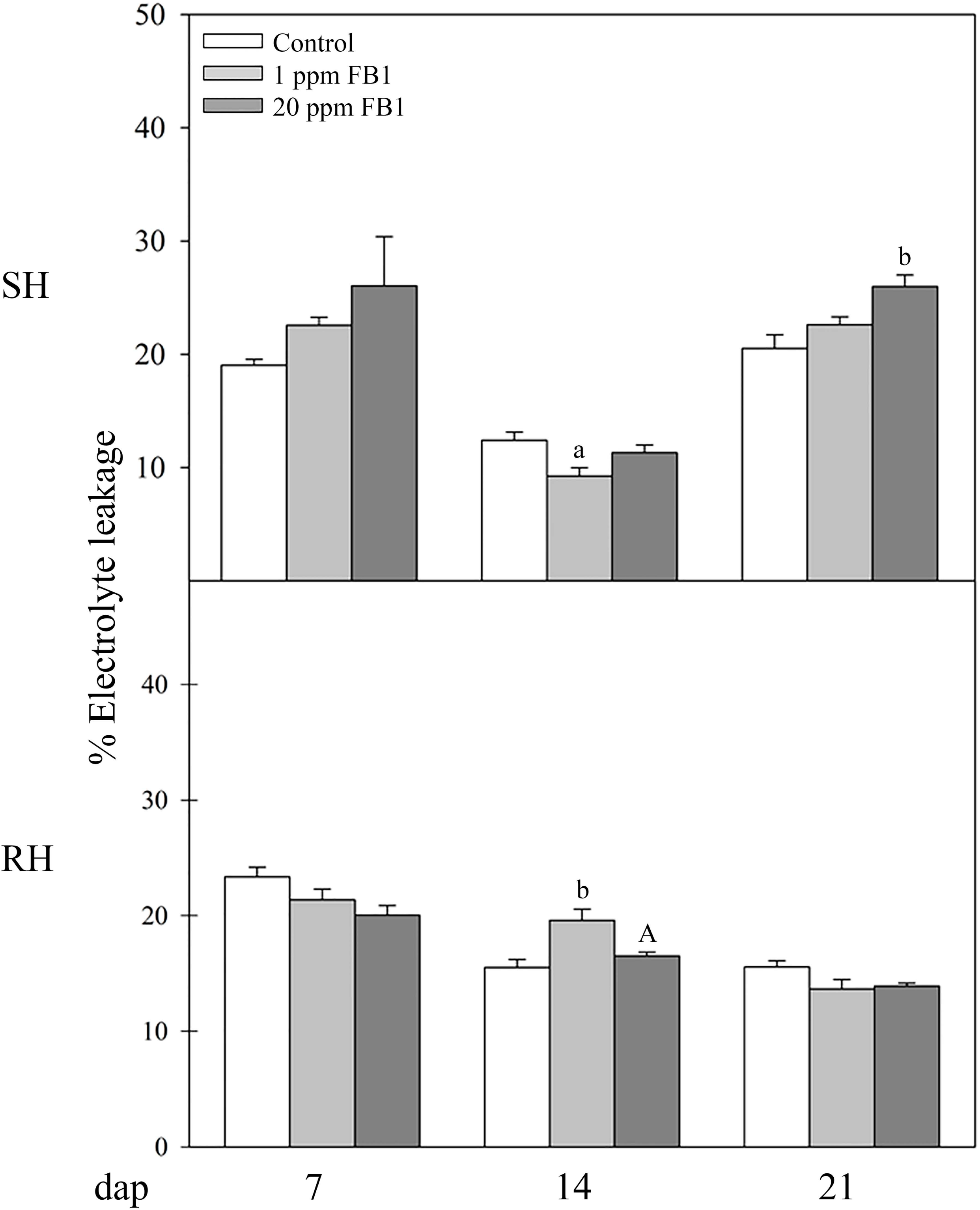
Effects of FB1 on electrolyte leakage in leafs of SH and RH maize seedlings. Data are represented as Mean ± SE. ^a/A^p < 0.05; ^b^p < 0.01. Lower case letters denote differences between treatments and control. Capital letters indicate differences between both FB1 exposure levels.

### Hydrogen peroxide

H_2_O_2_ was quantified in maize seedlings exposed or not to FB1. In general, little effects were observed in SH watered with the lowest toxin concentration (Fig. 2). H_2_O_2_ decreased at 7 dap in roots of both hybrids, whereas in aerial parts, a similar outcome was observed only in RH. Moreover, this ROS increased at 14 and 21 dap in aerial parts of RH, while, in roots, similar and lower H_2_O_2_ levels were found, respectively.

**Fig. 2.**
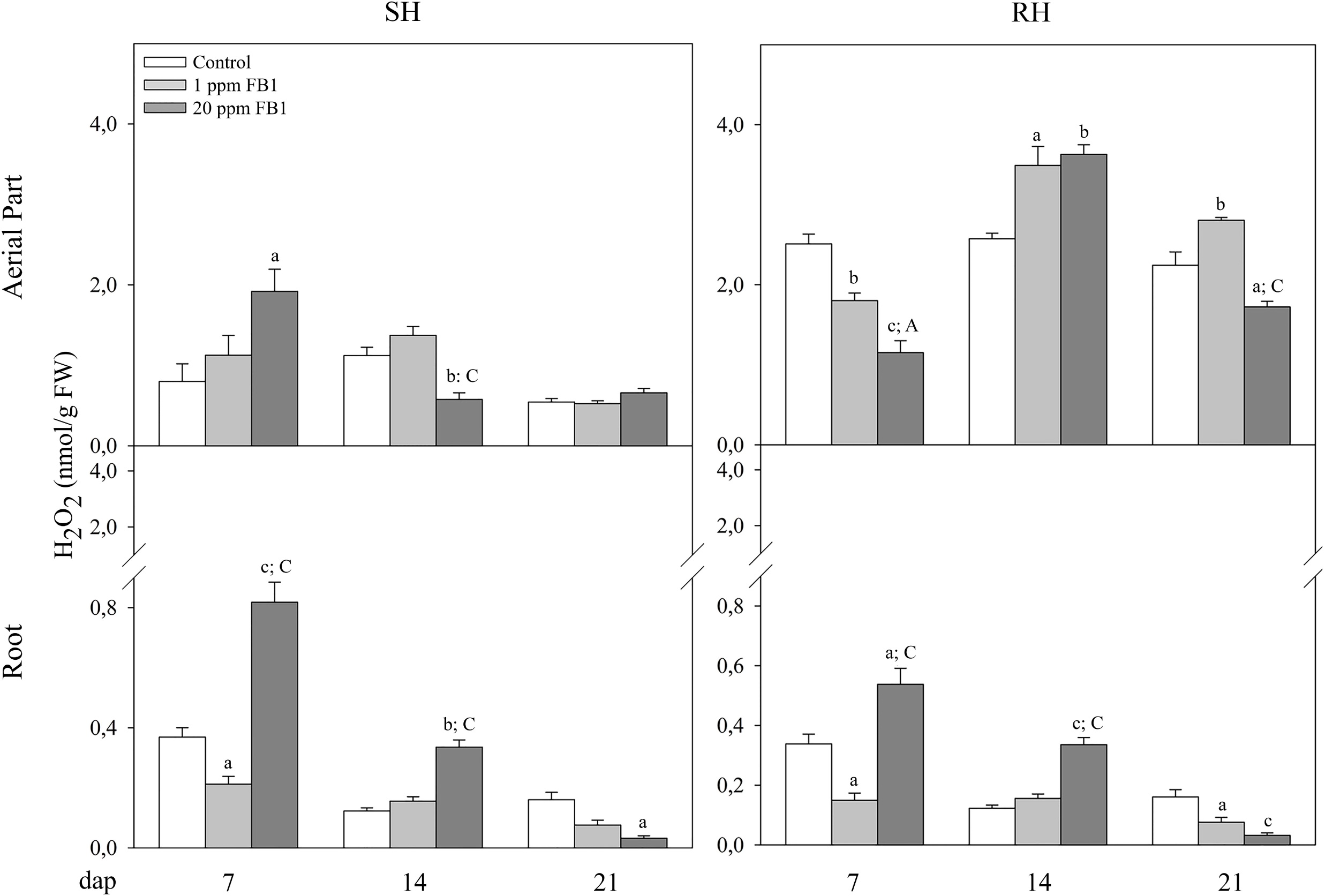
Effects of FB1 on hydrogen peroxide accumulation in aerial parts and roots of SH and RH maize seedlings. Data are represented as Means ± SE. ^a/A^p < 0.05; ^b^p < 0.01; ^c/C^p < 0.001. Lower case letters denote differences between treatments and control. Capital letters indicate differences between both FB1 exposure levels.

Watering with FB1 20 ppm increased the H_2_O_2_ in roots of both hybrids in almost all endpoints assessed, except for RH at 21 dap. Nevertheless, in aerial parts, such effects were only found at 7 dap in SH, and at 14 dap in RH.

### Antioxidant enzymes

The maize genotype susceptible to infection by *F. verticillioides* was characterised by higher basal SOD and GPOX activities compared with RH, which was evidenced in both roots and aerial parts, and in all endpoints assessed (Table 1). Furthermore, the effects of FB1 on these antioxidant activities were markedly different in both hybrids. In SH, the irrigation with 20 ppm of toxin at 7 dap increased GPOX enzymatic activities in roots, and SOD in aerial parts; while at 14 dap, the highest concentration of mycotoxin increased SOD activities in stems and leaves. However, the FB1 effects on this hybrid were mainly inhibitory of both enzymes. Minor SOD and GPOX activities were recorded at 7 dap in roots of seedlings irrigated with 1 ppm of FB1. Similar changes were caused by both toxin concentrations in roots at 14 and 21 dap and, in aerial parts, in the last endpoint assessed.

**Table 1:** Effects of FB1 on SOD and GPOX activities in aerial parts and roots of SH and RH maize seedlings ^1^.

| Enzyme | Plant part | FB1 (ppm) | SH |  |  | RH |  |  |
| --- | --- | --- | --- | --- | --- | --- | --- | --- |
|  |  |  | dap |  |  | dap |  |  |
|  |  |  | 7 | 14 | 21 | 7 | 14 | 21 |
| SOD | Aerial | 0 | 45.8±5.2 | 37.3±3.6 | 70.2±2.2 | 13.2±1.8 | 29.6±2.7 | 29.0±1.5 |
|  |  | 1 | 40.1±2.6 | 67.6±4.9<sup>c</sup> | 45.0±2.3<sup>c</sup> | 19.0±3.0 | 28.9±2.9 | 30.0±1.3 |
|  |  | 20 | 70.1±2.6<sup>b,C</sup> | 44.7±3.5<sup>B</sup> | 38.6±2.8<sup>c</sup> | 23.3±4.0<sup>a</sup> | 32.1±2.0 | 31.9±2.1 |
|  | Root | 0 | 56.7±3.4 | 83.3±14.8 | 141.4±26.3 | 18.5±2.5 | 44.0±1.4 | 45.8±3.6 |
|  |  | 1 | 40.4±2.4<sup>b</sup> | 43.8±2.9<sup>a</sup> | 41.2±2.3<sup>c</sup> | 25.2±5.4 | 34.7±3.5 | 38.9±3.1 |
|  |  | 20 | 60.4±3.6<sup>C</sup> | 39.4±2.2<sup>b</sup> | 34.9±4.5<sup>c</sup> | 45.2±2.5<sup>c,B</sup> | 69.5±4.6<sup>c,C</sup> | 50.8±9.3 |
| GPOX | Aerial | 0 | 147.0±17.3 | 56.5±8.1 | 95.9±5.8 | 20.1±0.8 | 26.3±1.0 | 42.9±1.3 |
|  |  | 1 | 136.4±9.5 | 63.6±6.5 | 74.9±0.9<sup>b</sup> | 36.7±1.7<sup>c</sup> | 26.7±1.0 | 36.0±1.0<sup>c</sup> |
|  |  | 20 | 115.4±5.5 | 57.9±8.2 | 58.3±0.8<sup>c,A</sup> | 30.5±1.3<sup>c,A</sup> | 40.2±2.5<sup>c,C</sup> | 34.6±0.3<sup>c</sup> |
|  | Root | 0 | 226.0±5.4 | 733.5±65.4 | 1536.6±165.7 | 126.7±3.6 | 474.2±6.8 | 330.6±23.1 |
|  |  | 1 | 194.0±9.4<sup>a</sup> | 553.3±10.6<sup>a</sup> | 365.5±13.7<sup>c</sup> | 188.0±8.3<sup>c</sup> | 507.9±21.6 | 422.7±5.4 |
|  |  | 20 | 264.2±10.8<sup>a,C</sup> | 405.3±21.4<sup>c,A</sup> | 289.8±14.4<sup>c</sup> | 148.3±4.9<sup>a,C</sup> | 304.9±6.4<sup>c,C</sup> | 529.4±45.1<sup>b</sup> |

<sup>1</sup> SOD (units/mg protein) and GPOX ( $\Delta\text{Abs}_{436 \text{ nm}} \cdot \text{mg protein}^{-1} \cdot \text{min}^{-1}$ ) activities are represented as Means ± SE. <sup>a/A</sup>p < 0.05; <sup>b/B</sup>p < 0.01; <sup>c/C</sup>p < 0.001. Lower case letters denote differences between treatments and Control. Capital letters indicate differences between both FB1 exposure levels. SH, susceptible hybrid; RH, resistant hybrid; dap, days after planting; SOD, superoxide dismutase; GPOX, guaiacol peroxidase.

Unlike the findings in SH, FB1 increased both antioxidant activities in RH, except for the GPOX decreases registered in roots at 14 dap, and at 21 dap in aerial parts of seedlings exposed to 20 and 1 ppm of toxin, respectively. Both toxin concentrations increased the GPOX throughout the seedlings at 7 dap; with 20 ppm, in aerial parts at 14 dap; and in both plant parts at 21 dap. In addition, the irrigation with 20 ppm of FB1 increased SOD throughout the seedlings at 7 dap, and in roots at 14 dap.

### TBARS

TBARS were measured in order to estimate the lipidic oxidative damages induced by FB1. TBARS were higher at 7 dap, and decreased at 21 dap in roots from both hybrid seedlings watered with 1 ppm of FB1 (Fig. 3). Despite these findings, TBARS increased in the aerial parts of the RH plantlets at 21 dap.

**Fig. 3.**
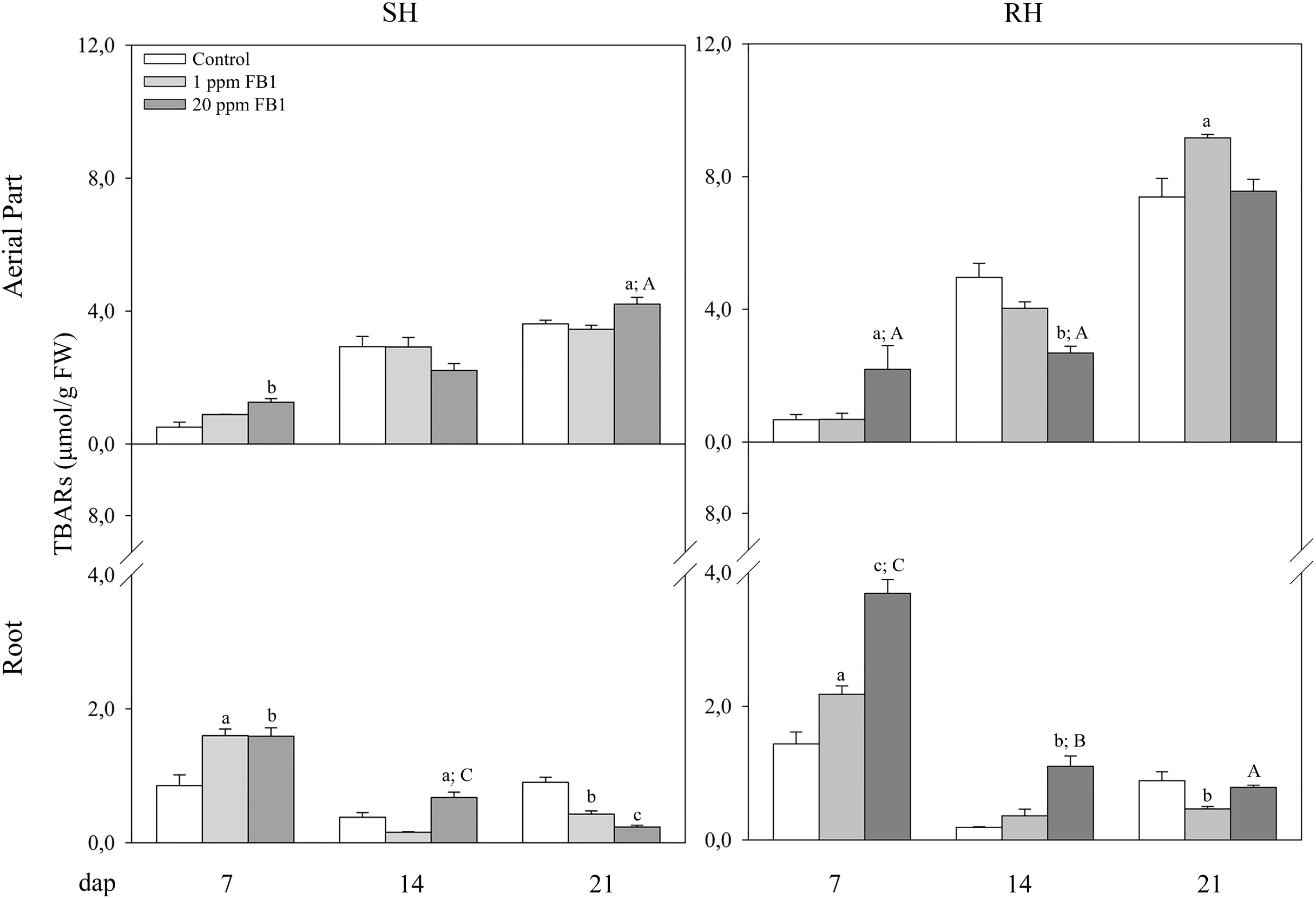
Effects of FB1 on TBARS accumulation in aerial parts and roots of SH and RH maize seedlings. Data are represented as Means ± SE. ^a/A^p < 0.05; ^b/B^p < 0.01; ^c/C^p < 0.001. Lower case letters denote differences between treatments and control. Capital letters indicate differences between both FB1 exposure levels.

Similar phytotoxic effects were observed in roots from SH and RH seedlings exposed to 20 ppm of FB1, where the mycotoxin raised the TBARS at 7 and 14 dap. However, a major lipidic oxidation in aerial parts was estimated in both hybrids at 7 (but not 14) dap, and at the last endpoint assessed in SH, where TBARS in roots decreased with respect to controls.

### Discriminant analysis and integrated biomarker response

An Integrated Biomarker Response index (IBR) was calculated with the aim to obtain a more complete understanding of biological effects suffered by the tested hybrids. In our study, the biomarkers selected through a discriminant analysis and used to calculate IBR values are informed in the Supplementary Material (Tables S1 and S2).

An integrated biomarker response (IBR) was then calculated for each treatment from the parameters selected by the discriminant analysis, allowing a comprehensive understanding of the stress level experienced by hybrids. The IBR calculated for every experimental condition is shown in Figure 4 and in Table 2. The grey areas shown in graphs, delimited by linking the IBR of control and FB1 (1 and 20 ppm) groups, allow a better visualisation of the treatment that produced the greatest stress. Both FB1 concentrations used in this study caused significant IBR increases with respect to control. Irrigation with 20 ppm of FB1 induced the greatest IBR in both hybrids, and in almost all the endpoints assessed, with the exception of RH at 21 dap, where the greatest stress was caused by the lowest concentration of the mycotoxin.

**Fig. 4.**
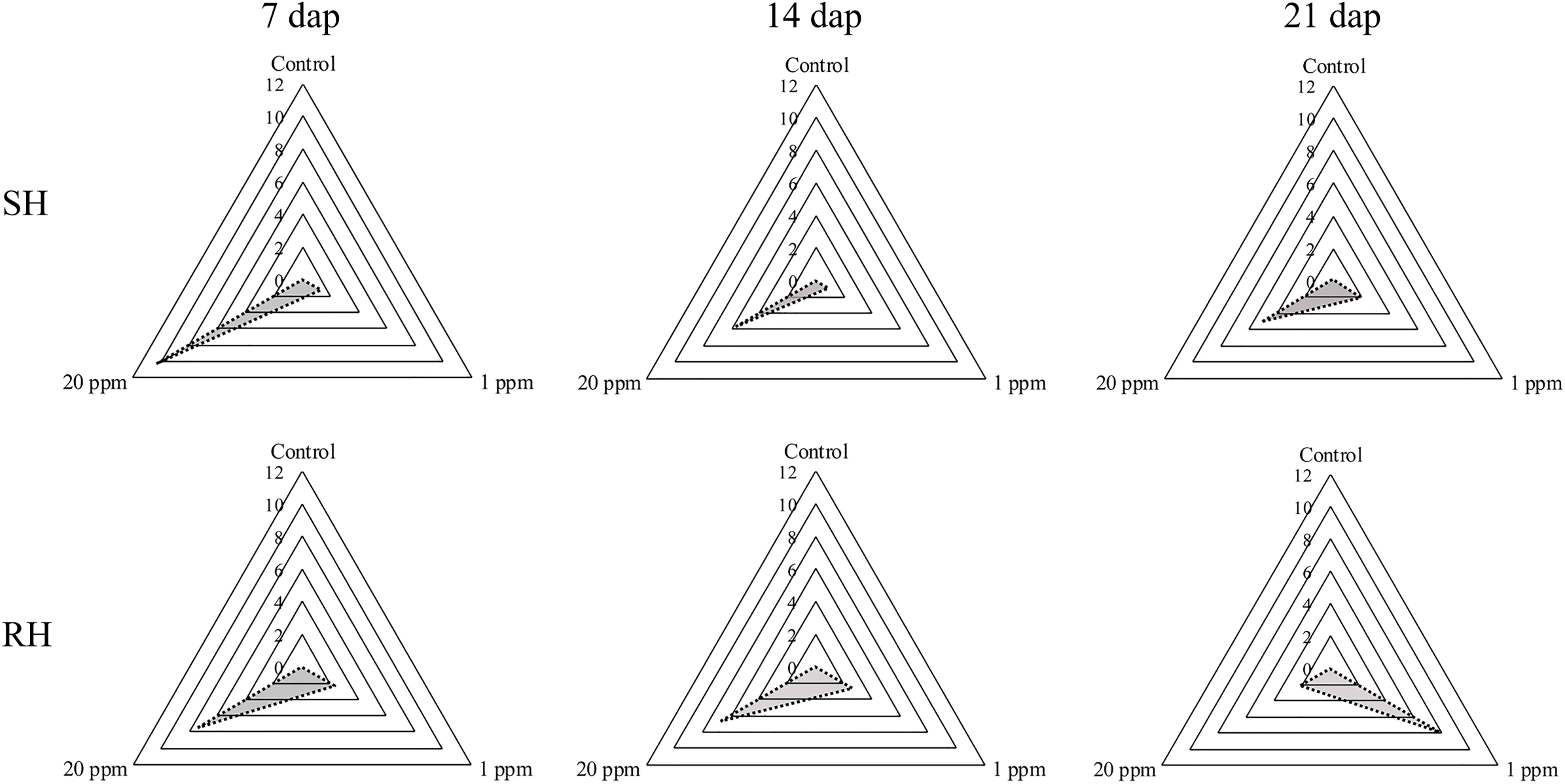
Integrated Biomarker Response (IBR) of SH and RH maize seedlings exposed at 0 (Control), 1 and 20 ppm of FB1 during 7, 14 and 21 days after planting (dap). Radar graph for the calculated IBR index. The spokes of the radar indicate the IBR index mean values for each studied treatment.

**Table 2.** Integrated biomarker response (IBR) in irrigations of SH and RH of maize at different concentrations of FB1.

|  |  |  | IBR |  |  |  |  |
| --- | --- | --- | --- | --- | --- | --- | --- |
|  | dap | FB1 (ppm) | Median |  | Media | Min | Max |
| SH | 7 | 0 | 0.00 |  | 0.07 | 0.00 | 0.18 |
|  |  | 1 | 1.29 | c | 1.37 | 0.82 | 1.96 |
|  |  | 20 | 10.31 | c, C | 10.31 | 10.30 | 10.32 |
|  | 14 | 0 | 0.00 |  | 0.00 | 0.00 | 0.00 |
|  |  | 1 | 0.85 | b | 0.85 | 0.02 | 1.68 |
|  |  | 20 | 5.72 | c, C | 5.72 | 5.64 | 5.81 |
|  | 21 | 0 | 0.13 |  | 0.13 | 0.00 | 0.26 |
|  |  | 1 | 1.96 | c | 1.96 | 1.15 | 2.78 |
|  |  | 20 | 5.04 | c, C | 5.04 | 4.96 | 5.12 |
| RH | 7 | 0 | 0.00 |  | 0.00 | 0.00 | 0.00 |
|  |  | 1 | 2.29 | c | 2.31 | 1.26 | 3.43 |
|  |  | 20 | 7.42 | c, C | 7.48 | 7.24 | 7.76 |
|  | 14 | 0 | 0.00 |  | 0.00 | 0.00 | 0.00 |
|  |  | 1 | 2.59 | c | 2.71 | 1.87 | 3.64 |
|  |  | 20 | 6.70 | c, C | 6.77 | 6.61 | 7.12 |
|  | 21 | 0 | 0.00 |  | 0.00 | 0.00 | 0.00 |
|  |  | 1 | 7.81 | c | 7.83 | 7.75 | 7.92 |
|  |  | 20 | 2.08 | c, C | 2.18 | 0.91 | 3.30 |

Median, mean, minimal and maximal values for each treatment. Different letters indicate Means
± SE. <sup>a/A</sup>p < 0.05; <sup>b/B</sup>p < 0.01; <sup>c</sup>p < 0.001. Lower case letters denote differences between treatments and Control. Capital letters indicate differences between both FB1 exposure levels.
SH, susceptible hybrid; RH, resistant hybrid; dap, days after planting.

### Mechanisms involved in the phytotoxicity of FB1: Hydroponic model Oxidative stress

We studied more deeply the mechanisms involved in the cell death caused by FB1 to maize seedlings. A hydroponic model was chosen for this purpose, due to the minimal interference of sample manipulation in the results. First, we assessed if the oxidative stress was associated with the phytotoxicity caused by FB1 (1 ppm) in the hydroponic model. The H_2_O_2_ content was evaluated in both hybrids exposed to FB1, with and without pre-treatment with ascorbic acid (AA), a widely used ROS scavenger. As shown in Fig. 5A, the pre-treatment of seedlings with AA prevented the H_2_O_2_ increase induced by FB1 in both hybrids, therefore confirming its antioxidant activity.

**Fig. 5.**
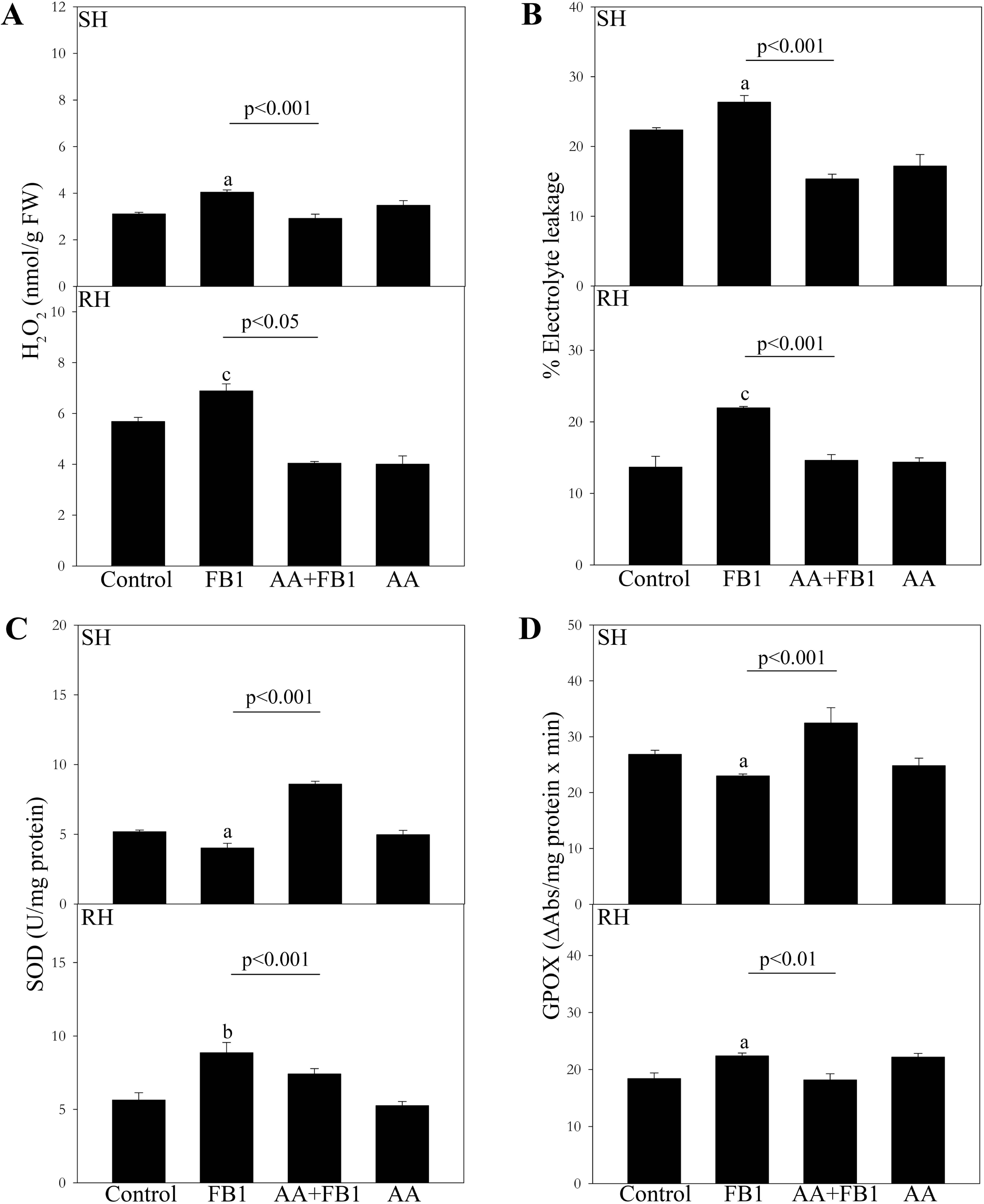
Effects of 1 ppm of FB1 on the content of A) hydrogen peroxide, B) electrolyte leakage, C) Superoxide dismutase, and C) guaiacol peroxidase in SH and RH maize seedlings pre-treated or not with 1 mM AA. Data are represented as Means ± SE. ^a^p < 0.05; ^b^p < 0.01; ^c^p < 0.001 indicate differences with the control. p-value indicated in the graph shows differences between FB1 vs AA + FB1 treatments.

Later, we studied if the cell death observed in seedlings grown in pots was also induced by FB1 in hydroponia, and its relation to the oxidative stress. The treatment with the mycotoxin increased the electrolyte leakage (EL) % at 24 hpt in both hybrids, but such outcomes were prevented by the pre-treatment of seedlings with AA (Fig. 5B). The consequences of the FB1 exposure in the SOD and GPOX antioxidant activities were similar to those found in seedlings grown in pots. While the mycotoxin decreased both activities in SH (Fig. 5C and D), the opposite was observed in RH. Nevertheless, such effects were prevented by the pre-treatment with the antioxidant.

### FB1-induced cell death: Modulatory effects of ROS on phytohormones

In order to explore the modulatory effects of ROS on phytohormones in the FB1-induced cell death, the levels of SA and JA in both hybrids were quantified. SA remained unaltered in SH seedlings treated with FB1, but JA was decreased (Fig. 6A and B). Moreover, despite the mycotoxin had no effects on SA, it increased the JA levels in the seedlings pre-treated with the antioxidant (with respect to those untreated with AA). In RH, the toxin had opposed effects on SA and JA levels (increase and decrease, respectively), but although the pre-treatment with AA prevented such SA rise, it could not prevent the fall of JA caused by the mycotoxin (Fig. 6A and B).

**Fig. 6.**
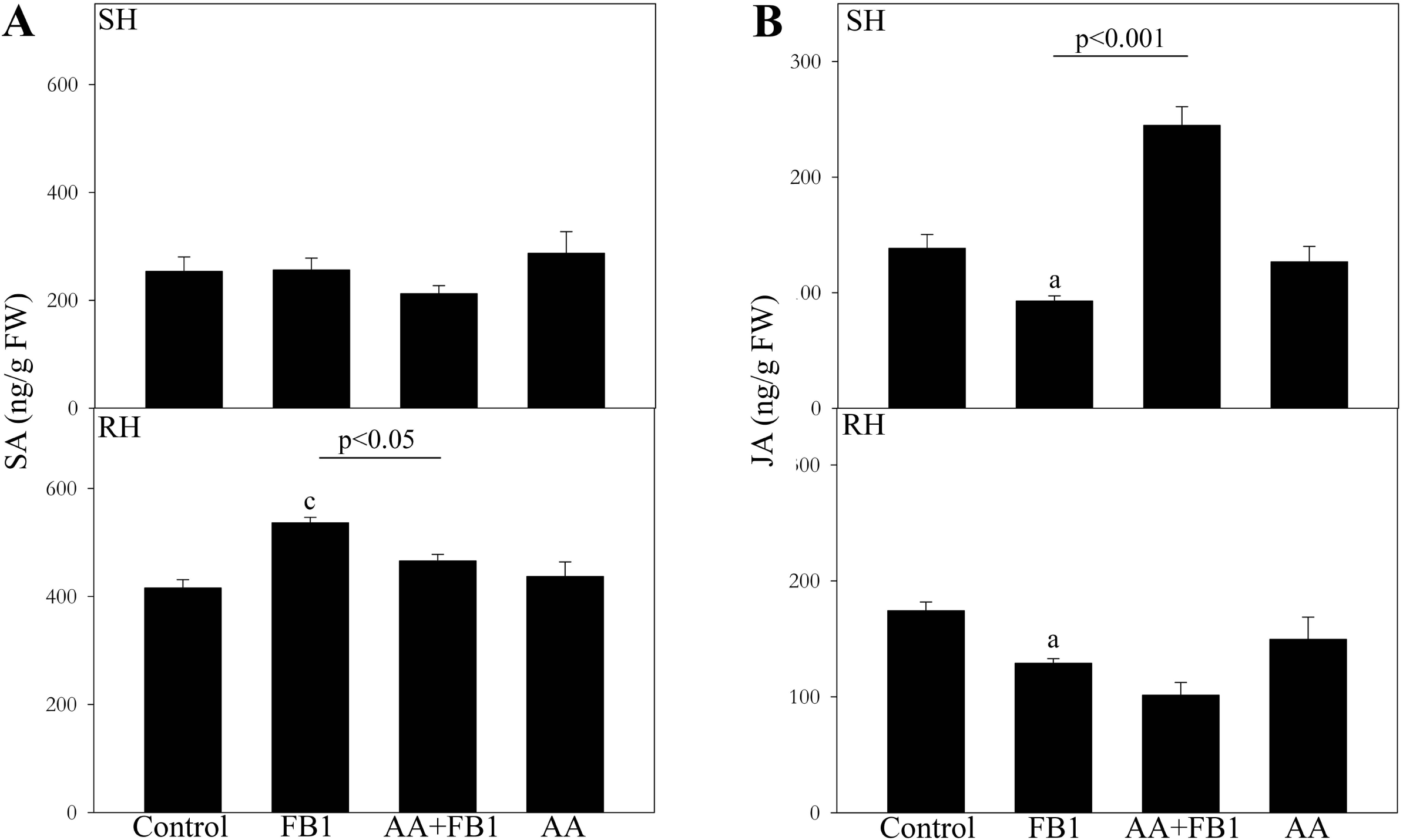
Effects of 1 ppm of FB1 on the content of SA and JA in SH and RH maize seedlings pre-treated or not with 1 mM AA. Means ± SE of the content of SA (A) and JA (A) are shown. ^a^p < 0.05; ^c^p < 0.001 indicate differences with the control. p-value indicated in the graph shows differences between FB1 vs AA + FB1 treatments.

## Discussion

The evidence collected to date shows FB1 as a phytotoxin apparently not essential for the pathogenicity of *F. verticillioides* in corn, although it may favour the fungal invasion of maize plant tissues. This mycotoxin is a potent inducer of programmed cell death in plants, and much of the progress in this field was in *Arabidopsis thaliana* as experimental model (Abbas *et al*., 1994; Stone *et al*., 2000; Xing *et al*., 2013; Glenz *et al*., 2019).

Having in mind that FB1 can be found in ground with corn debris (Abbas *et al*., 2008) and drainage water next to croplands (Waskiewicz *et al*., 2015), the toxin probably accumulates in soils and can debilitate maize seedlings growing on it, even in absence of fungal infection, since it can be absorbed from soil and disseminated throughout the plant to exert its toxicity (Arias *et al*., 2016). Therefore, we carried out two experimental designs: i) a “chronic phytotoxicity” model to characterise cell death and oxidative status in seedlings grown in pots up to 15 days after exposure to 1 and 20 ppm of FB1 (21 dap); and ii) an “acute phytotoxicity” model in hydroponics to assess the modulating effects of ROS on SA and JA, the phytohormones involved in the cell death induced at 24 h of treatment with 1 ppm of FB1. The exposure levels used in this work were chosen on the basis of previous works, where 1 and 20 ppm of FB1 reproduced the phenotype of corn seedlings infected by *F. verticillioides*, although plants could apparently detoxify 1 ppm of FB1 (Arias *et al*., 2016; Arias *et al*., 2012). Due to the higher biological relevance of this concentration, it was used for studying the FB1 acute phytotoxicity in maize.

Cell death may be provoked by mycotoxins as part of the fungal strategies to invade plants. In this study, we observed changes in electrolyte leakage (EL) which depended on the FB1 concentration, the hybrid, and the age of the plants. The highest EL induced at 14 dap in RH watered with 1 ppm of FB1 probably shows that the toxin caused the loss of the plasma membrane integrity, leading to higher ion permeability as the ultimate step in cell death. A similar result could have been observed at 21 dap in SH irrigated with 20 ppm of FB1. However, while in the first case (RH watered with 1 ppm of FB1) the EL increase was transient, we could not clarify whether this parameter returned to values comparable to the control after the last point assessed in SH. The meaning of the lowest EL observed at 14 dap in SH is also unclear. Taken together, these data show that cell death is a chronic toxic effect induced by FB1 in maize seedlings regardless of the hybrid susceptibility or resistance to *Fusarium* ear rot in the field, although the severity and the kinetics of its chronic phytotoxicity may depend on the host genetic background. Moreover, the EL registered here is a probable consequence of the differential sphinganine (Sa) and phytosphingosine (Pso) imbalances reported in maize seedlings from the same genotypes used in this work, upon FB1 exposure (Arias *et al*., 2016).

Maschietto and colleagues (2016) analysed two maize hybrids with contrasting resistance to *Fusarium* infection, and proposed that the resistant phenotype is related to the higher constitutive expression of antioxidant enzymes and defence-related proteins. However, the results of this work are not in line with this idea, since we observed greater constitutive SOD and GPOX activities in SH, highlighting the need to study a greater number of hybrids to reaffirm or discard an eventual direct relationship between resistance to *Fusarium* and high constitutive antioxidant activities. Also, it is important to note that the resistance to ear rot by *Fusarium* spp. is under polygenic control and strongly influenced by environmental factors (Presello *et al*., 2006).

We observed that the roots are the most affected plant parts when soils are contaminated by FB1. Further, considering the number and biological meaning of the alterations found in each condition, a major toxicity of the highest concentration of mycotoxin becomes evident. In SH, the changes were characterised by the inhibitory effects of the toxin (both concentrations) on the antioxidant enzymes, SOD and GPOX, as well as by the highest levels of H_2_O_2_ and TBARS induced by 20 ppm of FB1 in the three endpoints assessed. This toxin concentration also induced the major changes in RH, with H_2_O_2_ and TBARS increases, but unlike the findings from SH, higher SOD and GPOX activities were observed in RH. The phytotoxic effects of FB1 on the aerial parts of SH and RH, irrigated with both toxin concentrations, were less evident than those found in the roots, although they reflected the principal changes induced in the latter. Maschietto *et al*. (2016) observed that the activities of antioxidant enzymes increased more rapidly in a resistant genotype after inoculation with *F. verticillioides*. A similar behaviour was found in this work, where the increases in the SOD and GPOX activities were evident in the aerial parts and roots of RH seedlings watered with FB1, therefore suggesting that this mycotoxin could be involved in the faster enzymatic antioxidant response upon seedling infections by *F. verticillioides*. Moreover, the results of this work could show that, in the case of the resistant phenotype, the speed of the enzymatic antioxidant response of maize hybrids, rather than the basal enzymatic activities, would be more closely related to *F. verticillioides* ear rot in the field. However, it is important to emphasize that, apart from FB1, other soluble or structural fungal components could modulate the plant-fungus interactions. Also, several plant secondary metabolites, quantitatively less important than enzymes in the antioxidant defences, contribute to maintaining the redox balance (Bartoli *et al*., 2013; Noctor *et al*., 2018).

Plant growth is strongly influenced by external conditions, and the cellular redox homeostasis was proposed as a key biochemical connection between plant metabolism and environment (Foyer and Noctor, 2009; Noctor *et al*., 2018). In this sense, the integrated biomarker response (IBR) allowed us to get a comprehensive view of the stress produced by FB1 on the oxidative status of the plantlet cells. As it was expected, the stress evolved differently depending on the toxin concentration. The IBR pointed out that 1 ppm of FB1 was generally less stressful for both hybrids than the highest toxin concentration, with the exception of RH at 21 dap, where the opposite was observed. Using the same hybrids and experimental model, Arias *et al*. (2016; 2012) reported that the biomass and fitness of maize seedlings irrigated with 1 ppm of FB1 were restored at 21 dap, so it was proposed that they would have efficient detoxification / excretion mechanisms for this level of exposure to FB1. However, the toxin accumulation and the incidence and severity of lesions, both in aerial parts and roots, were greater in SH, showing different biochemical responses to this mycotoxin, depending on the maize genotype. The results of this work suggest that the antioxidant systems of RH would respond more efficiently to control the oxidative stress caused by FB1.

The IBR also evidenced two clearly different responses of both germplasms to irrigation with the lowest concentration of toxin. FB1 inhibited the antioxidant SOD and GPOX in SH, but here IBRs were only slightly higher, which could be suggesting that the remaining enzymatic activities would be enough to control the pro-oxidant effect of the toxin at all endpoints assessed. This interpretation is also supported by the overall behaviour of H_2_O_2_ and TBARs, biomarkers that in SH were mostly unaltered by the toxin. A different scenario could have been occurring in RH, where the transient increase in cell death found at 14 dap might be showing that FB1 induced hypersensitive response (HR)-type cell death, an ordered process probably triggered by the oxidative stress, that would allow to restrict the mycotoxin to a part of the plant (Stone *et al*., 2000; Xing *et al*., 2013). Such host response is not fundamental for the generation of resistance, but it is required for a rapid and strong activation of both local and systemic defence mechanisms (Heath, 2000). In this regard, the highest IBR found at 21 dap might be denoting a plant stress caused by its continuous response to alleviate the phytotoxicity of a contaminant that persists in soil.

High levels of antioxidant enzymes allow the plant to maintain the redox homeostasis by rapidly scavenging the excess of ROS, thus, diminishing their toxic effects (Caverzan *et al*., 2016). Nevertheless, the success of acclimation to chronic stress by FB1 would include a more complex universe of regulatory mechanisms of the cellular redox state, focused not only on the antioxidant system, but also on the signalling mediated by the ROS itself.

Beside generating cell damage in the plant, EROs can act as second messengers by activating or inhibiting SA- and JA-mediated response mechanisms, respectively (Kwak *et al*., 2006; Noctor, 2018). The hydroponic model allowed us to evaluate the participation of these phytohormones in the acute phytotoxicity of FB1, minimizing the effects of the collection and conditioning procedures of the samples. Alike the observations in pots, the mycotoxin also induced acute cell death in both hybrids. In RH, it was associated with an increase in SA and a reduction in JA levels, dependent or not on the accumulation of ROS, respectively. These results are in line with those reported by Xing *et al*. (2013), who observed that the pre-treatment of *Arabdopsis thaliana* leaves with ascorbic acid prevented the SA rise upon infiltration of the leaves with 10 µM (7.22 ppm) of FB1, suggesting that this phytohormone would be ROS-modulated. However, the cell death in SH was only related to a ROS-mediated decrease of JA. The toxin was phytotoxic to both hybrids, but the highest cell death observed in RH would be related to the SA increase mediated by ROS, taking into account that FB1 reduced JA levels in both cases, as a result of two different regulatory mechanisms.

In summary, in this work we showed that FB1 caused cell death by two different biochemical mechanisms in hybrids with contrasting susceptibility to *F. verticillioides* ear rot in the field. In addition, the results suggest that ROS has a dual role in the mycotoxin-induced cell death in maize plants, generating oxidative stress, and modulating phytohormone-mediated defence responses to reduce the phytotoxicity of FB1. The balance between acclimation and cell death responses after the first contact with this mycotoxin would determine the fate of the plant. RH had a more efficient control of the FB1phytotoxicity, but the integrated biomarker response (IBR) might be pointing out a major stress in this hybrid to mitigate the chronic stress caused by the toxin.

## Experimental

### Chemicals and reagents

Fumonisin B1 (FB1) analytical standard (purity > 95 %) was purchased from PROMEC (Programme on Mycotoxins and Experimental Carcinogenesis, Tygerberg, Republic of South Africa). A soluble fertiliser, with a composition of 15 % N [6.5 % nitrate, 8.5 % ammonia], 15 % P as P_2_O_5_, 15 % K as K_2_O and 3.2 % S was obtained from YARA (Buenos Aires, Argentina). Acetonitrile and methanol were of HPLC quality (Sintorgan, Argentina), and the other solvents used in this work were of analytical grade. 1,1,3,3-tetramethoxypropane (TEP, ≥ 97 %), 2-thiobarbituric acid (TBA, ≥ 98 %), superoxide dismutase (SOD), guaiacol peroxidase (GPOX) and trichloroacetic acid (TCA) were all purchased from Sigma-Aldrich, Buenos Aires, Argentina. The Bradford reagent was obtained from Bio-Rad Laboratories (Buenos Aires, Argentina). Ultrapure water (Millipore, Milli-Q system) was used to prepare standard solutions, dilutions and blanks.

### Fungal strain and inoculum preparation

A wild-type toxigenic isolate of *Fusarium verticillioides* (RC2024) obtained from carnation leaf-agar by monosporic isolation was used for fumonisins production. This strain was isolated from maize in Argentina, and stored in the Culture Collection Centre of the National University of Río Cuarto (RC), in Córdoba, Argentina. All cultures were maintained in 15 % glycerol at -80 °C. The ability of this strain to produce fumonisins was assessed using maize as the substrate, as previously described (Theumer *et al*., 2008). The RC2024 strain produced fumonisins at a ratio FB1:FB2:FB3 of 88:5:7.

Conidia suspensions were prepared with *F. verticillioides* RC2024 cultures grown at 25 °C for 7 days in V8 juice agar and Tween 20 at 2.5 % (v/v) in sterile water, and were used as inocula.

### Fumonisin production in bioreactor

The fumonisins used in the maize seedling assay were produced in liquid Myro medium as previously described by Arias *et al*. (2016; 2012). The fermentor vessel (10-L glass stirred-jar) (New Brunswick Scientific Co., Inc. Edison, NJ, USA) containing sterilised Myro medium ((NH_4_)_2_HPO_4_ (1 g), KH_2_PO_4_ (3 g), MgSO_4_.7H_2_O (2 g), NaCl (5 g), sucrose (40 g) and glycerin (10 g) in 10 L distilled-H_2_O) (Dantzer *et al*., 1996) was inoculated with the conidia suspension and maintained at 28 °C with 120 rpm agitation. Aerobic conditions were maintained using a stir rate and an air flow rate of 2 standard litres per minute. The pH was continually monitored during fermentation by a gel-filled pH probe, and maintained within the 3.5 ± 0.1 range by a controller which operates peristaltic pumps, assigned to perform 0.1 M H_3_PO_4_ or 0.1 M NaOH addition, and incubation was carried out for 28 days. The fermented liquid medium was autoclaved and then clarified through a 0.45 µm filter. A sample of the filtrate was used for fumonisin quantification.

### Fumonisin quantification in fermented Myro medium

HPLC with fluorescence detection was used to quantify fumonisins produced in bioreactor. Samples of the fermented Myro medium were diluted with CH_3_CN at a 1:1 ratio, and the quantification of the diluted extracts was performed following a methodology proposed by Shephard *et al*. (1990). An aliquot (50 µL) of the diluted samples was derivatised with o-phthaldialdehyde (200 µL) soln., obtained by adding 0.1 M sodium tetraborate (5 mL) and 2-mercaptoethanol (50 µL) to MeOH (1 mL) containing o-phthaldialdehyde (40 mg). The derivatised samples were analysed by a Hewlett Packard series 1100 HPLC system, with a loop of 20 µL, and an isocratic pump (G1310A) coupled with a fluorescence detector (Agilent Technologies series 1200), at wavelengths of 335 nm and 440 nm for excitation and emission, respectively. The column used was a 150 x 4.6 mm, 5 µm, Luna 100 RP-18, with a guard column of the same material (Phenomenex, Torrance, CA, USA). The mobile phase was MeOH-0.1M NaH_2_PO_4_ (75:25), with the pH being set at 3.35 ± 0.20 with o-phosphoric acid, and a flow rate of 1.5 mL/min was used. The quantitation of fumonisins was carried out by comparing the peak areas obtained from samples with those corresponding to analytical standards of FB1, FB2 and FB3 (purity > 95 %), using an HP Chemstation Rev. A.07.01 software.

## Maize seedling assays

### Phytotoxicity of FB1 in maize seedlings grown in pots

The maize (*Z. mays* L.) seedlings were obtained by sowing seeds of a resistant hybrid (RH; LT 622 MG) and a susceptible hybrid (SH; HX 31P77), which have shown resistance and susceptibility to *Fusarium* ear rot in the field, respectively (Presello *et al*., 2009).

The maize seeds were surface-disinfected for 2 min in 10 % bleach (0.4 % NaClO), rinsed three times with sterile H_2_O, and blotted dry on paper towelling. Then, seeds (three replicates of 10 seeds each) were sown in 24-cm diameter pots containing washed autoclaved sand, thus mimicking the simplest soil system with very little organic material or mineral nutrients (Arias *et al*., 2016; Arias *et al*., 2012). A soluble fertiliser was applied before planting and also twice a week thereafter. Pots were watered with FB1 solutions (1 and 20 ppm in sterile H_2_O, 100 mL) on days 2, 4, and 6 after planting, and then watered every 3 days with sterile water. The plants were grown under controlled conditions in a greenhouse with a 14/10 h light/dark cycle at 22 °C, and harvested 7, 14 and 21 days after planting (dap). Maize seedlings from all endpoints were collected for measuring electrolyte conductivity, H_2_O_2_, antioxidants enzymes (superoxide dismutase, SOD; and guaiacol peroxidase, GPOX), thiobarbituric acid reactive substances (TBARS), and chlorophylls quantification. Upon harvesting, leaf discs were immediately obtained from some seedlings (n=6 per group) for electrolyte conductivity measuring. The remaining seedlings were gently washed, and the roots were separated from the aerial parts of the plants. Both roots and aerial parts were ground to a powder after freezing with liquid N_2_ and kept at -80 °C until use.

### Mechanisms involved in the phytotoxicity of FB1: Hydroponic model

The maize seeds were surface-disinfected as described above. For hydroponic cultures, SH and RH maize seeds were submerged in a 1 mM CuSO_4_ solution at 25 °C for 24 hours. Then, they were incubated for 3 additional days between filter paper layers moistened with the same solution. Subsequently, the germinated seeds were transferred to 15 mL Falcon tubes (one per tube), containing hydroponic solution (0.25X). The concentration of this solution was gradually increased to 0.5X and 1X after 2 and 4 days of hydroponic culture, respectively. A hydroponic solution was used (2.5 mM Ca(NO_3_)_2_, 1.0 mM K_2_SO_4_, 0.2 mM KH_2_PO_4_, 0.6 mM MgSO_4_, 5.0 mM CaCl_2_, 1.0 mM NaCl, 1.0 µM H_3_BO_4_, 2.0 µM MnSO_4_, 0.5 µM ZnSO_4_, 0.3 µM CuSO_4_, 0.005 µM (NH_4_)_6_Mo_7_O_24_, 200 µM Fe–EDTA) (Zörb *et al*., 2013), which was changed every two days to avoid total consumption of nutrients. After 14 days, the aerial part of the seedlings was sprayed with 0 and 1 mM ascorbic acid (AA), 2 hours before the mycotoxin treatment (Xing *et al*., 2013). Then, the seedlings were exposed to 0 (Control) and 1 ppm of FB1 (dissolved in hydroponic solution). They were harvested at 24 hours post-treatment (hpt) with the mycotoxin, conditioned and stored as described above.

### Electrolyte conductivity

Cell death was assayed by measuring electrolyte leakage (EL) from leaf discs as described by Rizhsky *et al*. (2002), with minor modifications. Briefly, six leaf discs (6-mm diameter) were floated on 10 mL of ultrapure water and shaken at 60 rpm for 2 h at room temperature. Following incubation, the conductivity of the bathing solution was measured with a conductivity meter (CD 4301, Lutron). The solutions were then boiled at 95°C for 25 min to completely lyse the plant cell walls. The electrolyte conductivities of boiled solutions were recorded as the absolute conductivity. The percentage of EL was calculated as the initial conductivity / absolute conductivity x 100.

### Hydrogen peroxide

Hydrogen peroxide was measured spectrophotometrically following a procedure published by Alexieva *et al*. (2001). Ground tissues (0.3 g) were homogenised with 0.1 % trichloroacetic acid (1.5 mL), and then centrifuged (12,000 x g for 15 minutes at 4 °C). The reaction mixture consisted of 160 μL of 0.1 % TCA tissue extract supernatant, 160 µL of 100 mM KH_2_PO_4_/K_2_HPO_4_ buffer (pH 6.8) and 680 µL of 1 M KI solution in distilled water. Trichloroacetic acid (0.1 %) in absence of tissue extract was used as blank. The reaction was developed for 1 h in darkness, and absorbance measured at 390 nm using a microplate reader (Bio-Tek, Synergy HT). The amounts of H_2_O_2_ in samples were calculated using a standard curve (range: 0 – 1 mM), and the results were expressed as µmol (Fig 2) H_2_O_2_/g fresh weight (FW).

### Enzyme extraction and measurement

Enzyme extracts were prepared from individual plants according to Monferrán *et al*. (2009), with minor modifications. Ground tissues were homogenised with rupture buffer containing 0.1 M Na_2_HPO_4_/NaH_2_PO_4_ pH 6.5, 20 % glycerol, 1 mM EDTA, and 1.4 mM dithioerythritol. After removal of cell debris (10 min at 13,000 g), the supernatant was used for protein (Bradford, 1976) and enzyme measurements, which were determined by spectrophotometry using a microplate reader (Bio-Tek, Synergy HT).

The SOD activity was determined in 96 well plates according to the procedure described by Aiassa *et al*. (2010). Under illumination, riboflavin loses an electron and induces superoxide anion radical (O_2_^•-^), which reduces the nitroblue tetrazolium (NBT), but this last step was prevented by the SOD activity. The reaction mixture consisted of 10 µL of protein extract, SOD standard (calibration curve) or rupture buffer (blank); 30 µL of methionine 47.7 mM, 10 µL of NBT 0.825 mM in PBS, 30 µL of EDTA 0.367 µM and 30 µL of riboflavin 7.33 µM. The microplate was exposed to 20W fluorescent light for 30 minutes, and the colour developed was spectrophotometrically measured at 595 nm. The SOD activities in samples were expressed in units/mg protein, extrapolating the readings from samples in a calibration curve made with an analytical standard of SOD (0.25-1.00 μg/mL, equivalent to 1.14-2.56 SOD units/mL).

The GPOX activity was determined using H_2_O_2_ and guaiacol according to a procedure previously described (Bertrand *et al*., 2016). Briefly, 180 μL of Na_2_HPO_4_/NaH_2_PO_4_ (0.1M, pH 5.0) were mixed with 8.5 μL of guaiacol (100 mM in DMSO) and 8.0 μL of H_2_O_2_ (200 mM in DMSO). Then 10 μL of protein extract or rupture buffer (blank) were added, and the reaction mixture was incubated at 37 °C. Absorbances (436 nm) were recorded up to 4 minutes of reaction. The GPOX activity was expressed as the ΔAbs 436 nm.mg protein^-1^.min^-1^.

### Thiobarbituric acid reactive substances

TBARS were determined as indicators of lipid peroxidation according to a methodology proposed by Heath and Packer (1968), with minor modifications. Briefly, 0.5 g of ground tissue (aerial parts and roots) was homogenised with 2.5 mL of TCA 20 % (w/v) and centrifuged at 12,000 g for 4 minutes at 4 °C. Equal volumes of supernatant and reagent (thiobarbituric acid, TBA, 0.5 % dissolved in TCA 20 %) were then mixed. The samples were heated at 95 °C for 25 minutes, cooled in an ice bath, and then centrifuged at 9,000 g for 6 minutes at 4 °C. The absorbance at 532 nm was measured in the supernatant against a TBA blank, subtracting the absorbance of turbidity at 600 nm. The amounts of TBARS were calculated from a calibration curve based on the acid hydrolysis of TEP (0-100 µM) and the reaction with TBA, and the results were expressed as nmol TBARS/g of fresh weight tissue.

### Quantification of phytohormones by LC-MS/MS

The levels of jasmonic acid (JA) and salicylic acid (SA) in the aerial portion of plants were quantified. The extraction was carried out according to the method of Pan *et al*. (2008), with some modifications. Briefly, 0.5-1.0 g of tissue previously pulverised with liquid N_2_ were weighed, homogenised with 500 μL of 1-propanol/H_2_O/concentrated HCl (2:1:0.002; v/v/v), and stirred for 30 minutes at 4 °C. Then 1 ml of dichloromethane (CH_2_Cl_2_) was applied, stirred for 30 min at 4 °C, and centrifuged at 13000 g for 5 min. The lower organic phase (approx. 1 mL) was collected in vials, which was evaporated in a gaseous N_2_ sequence. Finally, it was re-dissolved with 0.1 - 0.15 mL of 100 % methanol (HPLC grade), and stirred slightly with vortex.

The system of the 1200 series of Agilent technologies (Agilent Technologies, Santa Clara, CA, USA) is equipped with a gradient pump (Agilent G1312B SL Binary), solvent degasser (Agilent G1379 B), auto sampler (Agilent G1367 D SL+WP) and a reversed phase column (C18 kinetex 2,6 µm, 100 mm x 2,1 mm, Phenomenex, Torrance, CA, USA). It is used as a mobile solvent system composed of water with 0.1 % HCO_2_H (A) and MeOH with 0.1% HCO_2_H (B), with a correction flow of 0.25 mL/min. The initial gradient of B was maintained at 30% for 2 min, and then linearly increased to 100% at 28 min. For identification and quantification purposes, a mass spectrometer of the microTOF-Q11 Series QTOF (Bruker, Billerica, MA, USA) coupled to the above mentioned HPLC (LC-MS / MS) was used. The ionization source was used with electrospray (ESI) and the Compass (version 3.1) and Data Analysis (version 4.1) programs were used for data acquisition and processing, respectively. The mass spectra of the data are recorded in negative mode. The mass/charge ratio (m/z) for each metabolite were: SA: 137.02 and JA: 209.12. The quantification of the activity was done by respecting the calibration curves with the linear adjustment, obtaining the results in a nanogram phytohormone/gram of fresh weight.

## Data analysis

### Integrated biomarker response

In order to achieve a more complete understanding of the seedlings reactions to treatments, an integrated biomarker response (IBR) was calculated with the aim to identify the level of response or stress expressed by the exposed organisms. In our study, those biomarkers (from aerial part and root) with greater ability to segregate tested conditions were selected by a discriminant analysis (forward method) using the Statistica Software (version 8.0).

This IBR was performed according to Beliaeff and Burgeot (2002), with modifications by Devin *et al*. (2014). Several IBRs were calculated, using a R Studio software (version 0.99.902), from the same data changing the order of the biomarkers. The final index value for each treatment was the calculated median. Finally, a Kruskall Wallis test was carried out to identify IBR differences between treatments.

### Statistical evaluation

Data from the toxicity studies were analysed by a two-tailed ANOVA, followed by a *post hoc* test (Bonferroni Multiple Comparisons) when the data presented homoscedasticity. In some cases, due to a lack of homoscedasticity, a nonparametric comparison was also performed using the Kruskal–Wallis test (p < 0.05). Differences were considered to be statistically significant for p values ˂ 0.05. The GraphPad InStat software version 3.01 (La Jolla, CA 92037 USA) was used for the analyses.

## Authors’ contributions

MGT conceived and designed research. SNOG conducted experiments. SLA, VSM, LB, PAV, MGR y HRR contributed to conduct experiments and analyse data. SNOG and MGT wrote the manuscript. All authors have read and approved the manuscript.

## Acknowledgements

This study was supported by grants from Secretaría de Ciencia y Tecnología-Universidad Nacional de Córdoba; Agencia Nacional de Promoción Científica y Tecnológica (PICT 2012-1742, 2013-0750 and 2015-2810). MGR hold fellowship from Agencia Nacional de Promoción Científica y Tecnológica. SNOG, PAV and LB hold fellowships from the Consejo Nacional de Investigaciones Científicas y Técnicas (CONICET-Argentina). MGT and VSM are career investigators of the latter institution.

The content of this work is the sole responsibility of their authors and does not necessarily represent the official views of the organisms that funded this research.

## Abbreviations

CAT: catalase; dap, days after planting
EL: electrolyte leakage
FB1: fumonisin B1
GPOX: guaiacol peroxidase
MDA: malondialdehyde
O_2_^•-^: superoxide radical anion
RH: resistant hybrid
ROS: reactive oxygen species
SH: susceptible hybrid
SOD: superoxide dismutase
TBA: thiobarbituric acid
TBARS: thiobarbituric acid reactive substances
TCA: trichloroacetic acid; SA, salicylic acid
JA: jasmonic acid
AA: ascorbic acid

## Supplementary Material

**Supplementary Table S1.** Discriminant analysis performed to identify those biomarkers measured in the susceptible hybrid (SH) exposed to 0; 1 and 20 ppm of FB1 during 7, 14 and 21 dap; and with higher contribution to discriminate among groups (exposure concentrations). Analyses were carried out with Statistica 8.0 software and biomarkers for each treatment were selected using a forward method. The classification matrix indicated that the model fitted led to an error of 10% with 100% correct assignation for each exposure concentration.

| **7 dap** |  |  |
| --- | --- | --- |
| Step 7, N of vars in model: 7; Grouping: Trat (3 grps) |  |  |
| Wilks' Lambda: ,00012 approx. F (14,36)=230,04 |  |  |
| p<0,0000 |  |  |
| **Biomarker** | **Wilks**<br>**Lambda** | **Parcial**<br>**Lambda** |
| H<sub>2</sub>O<sub>2</sub> <sub>root</sub> | 0,001026 | 0,119105 |
| TBARS <sub>root</sub> | 0,001649 | 0,074094 |
| TBARS <sub>aerial part</sub> | 0,000302 | 0,404607 |
| H<sub>2</sub>O<sub>2</sub> <sub>aerial part</sub> | 0,000224 | 0,544817 |
| EL | 0,000184 | 0,663963 |
| GPOX <sub>root</sub> | 0,000141 | 0,863800 |

| **14 dap** |  |  |
| --- | --- | --- |
| Step 5, N of vars in model: 5; Grouping: Trat (3 grps) |  |  |
| Wilks' Lambda: ,00129 approx. F (10,40)=107,29 |  |  |
| p<0,0000 |  |  |
| **Biomarker** | **Wilks**<br>**Lambda** | **Parcial**<br>**Lambda** |
| H<sub>2</sub>O<sub>2</sub> <sub>root</sub> | 0,004212 | 0,306699 |
| TBARS <sub>aerial part</sub> | 0,003377 | 0,382493 |
| TBARS <sub>root</sub> | 0,001781 | 0,725490 |
| EL | 0,001486 | 0,869109 |

| **21 dap** |  |  |
| --- | --- | --- |
| Step 5, N of vars in model: 5; Grouping: Trat (3 grps) |  |  |
| Wilks' Lambda: ,00218 approx. F (10,40)=81,733 |  |  |
| p<0,0000 |  |  |
| **Biomarker** | **Wilks**<br>**Lambda** | **Parcial**<br>**Lambda** |
| TBARS <sub>root</sub> | 0,011201 | 0,194345 |
| TBARS <sub>aerial part</sub> | 0,005470 | 0,397937 |
| SOD <sub>aerial part</sub> | 0,004750 | 0,458277 |
| H<sub>2</sub>O<sub>2</sub> <sub>root</sub> | 0,002830 | 0,769077 |

**Supplementary Table S2.**
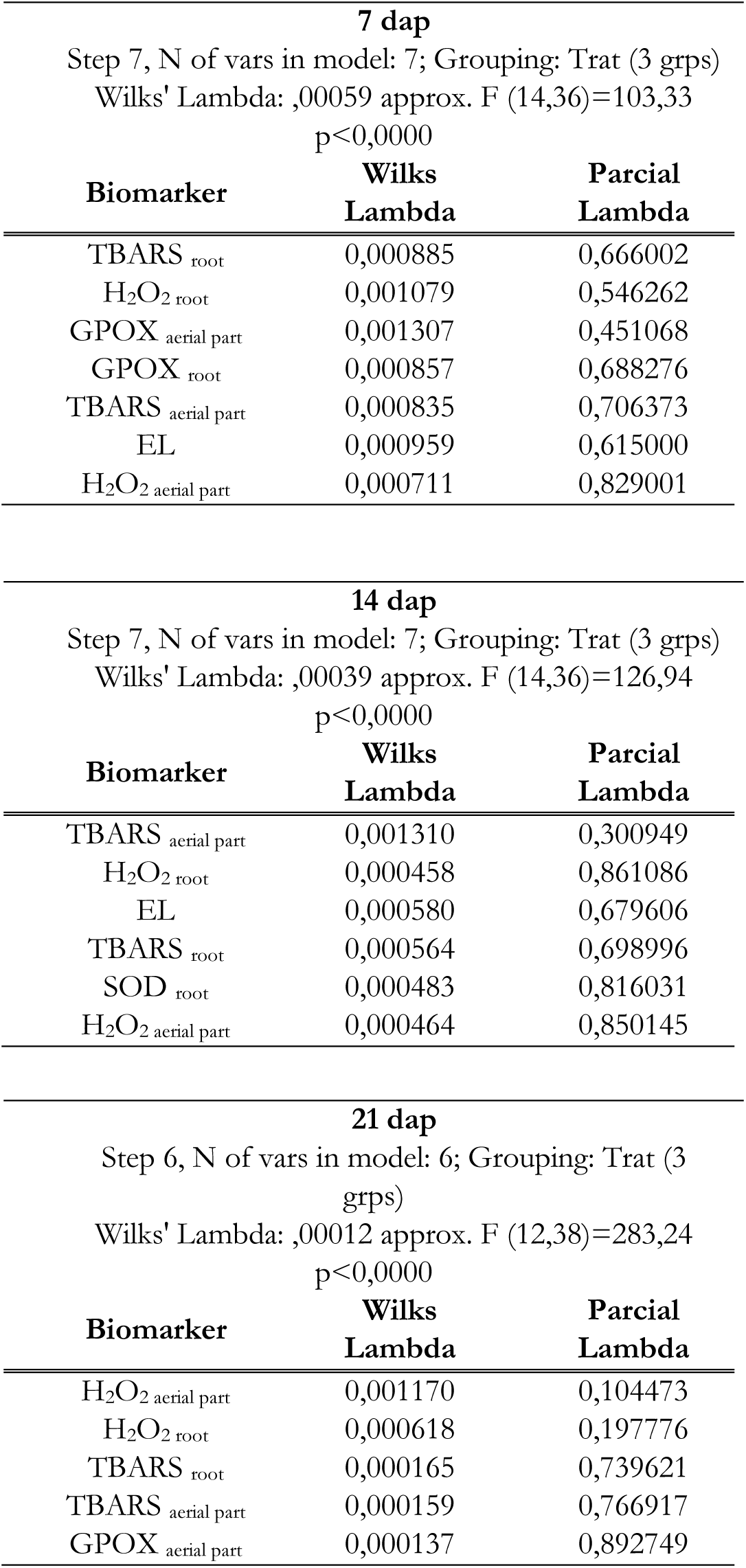
Discriminant analysis performed to identify those biomarkers measured in the resistant hybrid (RH) exposed to 0; 1 and 20 ppm of FB1 during 7, 14 and 21 dap; and with higher contribution to discriminate among groups (exposure concentrations). Analyses were carried out with Statistica 8.0 software and biomarkers for each treatment were selected using a forward method. The classification matrix indicated that the model fitted led to an error of 10% with 100% correct assignation for each exposure concentration.

| **7 dap** |  |  |
| --- | --- | --- |
| Step 7, N of vars in model: 7; Grouping: Trat (3 grps) |  |  |
| Wilks' Lambda: ,00059 approx. F (14,36)=103,33 |  |  |
| p<0,0000 |  |  |
| **Biomarker** | **Wilks**<br>**Lambda** | **Parcial**<br>**Lambda** |
| TBARS<sub>root</sub> | 0,000885 | 0,666002 |
| H<sub>2</sub>O<sub>2</sub><sub>root</sub> | 0,001079 | 0,546262 |
| GPOX<sub>aerial part</sub> | 0,001307 | 0,451068 |
| GPOX<sub>root</sub> | 0,000857 | 0,688276 |
| TBARS<sub>aerial part</sub> | 0,000835 | 0,706373 |
| EL | 0,000959 | 0,615000 |
| H<sub>2</sub>O<sub>2</sub><sub>aerial part</sub> | 0,000711 | 0,829001 |

| **14 dap** |  |  |
| --- | --- | --- |
| Step 7, N of vars in model: 7; Grouping: Trat (3 grps) |  |  |
| Wilks' Lambda: ,00039 approx. F (14,36)=126,94 |  |  |
| p<0,0000 |  |  |
| **Biomarker** | **Wilks**<br>**Lambda** | **Parcial**<br>**Lambda** |
| TBARS<sub>aerial part</sub> | 0,001310 | 0,300949 |
| H<sub>2</sub>O<sub>2</sub><sub>root</sub> | 0,000458 | 0,861086 |
| EL | 0,000580 | 0,679606 |
| TBARS<sub>root</sub> | 0,000564 | 0,698996 |
| SOD<sub>root</sub> | 0,000483 | 0,816031 |
| H<sub>2</sub>O<sub>2</sub><sub>aerial part</sub> | 0,000464 | 0,850145 |

| **21 dap** |  |  |
| --- | --- | --- |
| Step 6, N of vars in model: 6; Grouping: Trat (3 grps) |  |  |
| Wilks' Lambda: ,00012 approx. F (12,38)=283,24 |  |  |
| p<0,0000 |  |  |
| **Biomarker** | **Wilks**<br>**Lambda** | **Parcial**<br>**Lambda** |
| H<sub>2</sub>O<sub>2</sub><sub>aerial part</sub> | 0,001170 | 0,104473 |
| H<sub>2</sub>O<sub>2</sub><sub>root</sub> | 0,000618 | 0,197776 |
| TBARS<sub>root</sub> | 0,000165 | 0,739621 |
| TBARS<sub>aerial part</sub> | 0,000159 | 0,766917 |
| GPOX<sub>aerial part</sub> | 0,000137 | 0,892749 |

## References

Abbas, H.K., Accinelli, C., Zablotowicz, R.M., Abel, C.A., Bruns, H.A., Dong, Y., Shier, W.T., 2008. Dynamics of mycotoxin and Aspergillus flavus levels in aging Bt and non-Bt corn residues under Mississippi no-till conditions. J Agric Food Chem 56, 7578–7585.

Abbas, H.K., Tanaka, T., Duke, S.O., Porter, J.K., Wray, E.M., Hodges, L., Sessions, A.E., Wang, E., Merrill, A.H., Jr., Riley, R.T., 1994. Fumonisin- and AAL-Toxin-Induced Disruption of Sphingolipid Metabolism with Accumulation of Free Sphingoid Bases. Plant Physiol. 106, 1085–1093.

Aiassa, V., Barnes, A.I., Albesa, I., 2010. Resistance to ciprofloxacin by enhancement of antioxidant defenses in biofilm and planktonic Proteus mirabilis. Biochem Biophys Res Commun 393, 84–88.

Alexieva, V., Sergiev, I., Mapelli, S., Karanov, E., 2001. The effect of drought and ultraviolet radiation on growth and stress markers in pea and wheat. Plant, Cell Environ. 24, 1337–1344.

Arias, S.L., Mary, V.S., Otaiza, S.N., Wunderlin, D.A., Rubinstein, H.R., Theumer, M.G., 2016. Toxin distribution and sphingoid base imbalances in Fusarium verticillioides-infected and fumonisin B1-watered maize seedlings. Phytochemistry 125, 54–64.

Arias, S.L., Theumer, M.G., Mary, V.S., Rubinstein, H.R., 2012. Fumonisins: probable role as effectors in the complex interaction of susceptible and resistant maize hybrids and Fusarium verticillioides. J. Agric. Food. Chem. 60, 5667–5675.

Asai T., Stone J.M., Heard J.E., Kovtun Y., Yorgey P., Sheen J., and Ausubel F.M. 2000. Fumonisin B1-induced cell death in arabidopsis protoplasts requires jasmonate-, ethylene-, and salicylate-dependent signaling pathways. Plant Cell 12, 1823–1836.

Bartoli, C.G., Casalongué, C.A., Simontacchi, M., Marquez-Garcia, B., Foyer, C.H., 2013. Interactions between hormone and redox signalling pathways in the control of growth and cross tolerance to stress. Environ. Exp. Bot. 94, 73–88.

Beliaeff, B., Burgeot, T., 2002. Integrated biomarker response: a useful tool for ecological risk assessment. Environ Toxicol Chem 21, 1316–1322.

Bertrand, L., Asis, R., Monferrán, M.V., Amé, M.V., 2016. Bioaccumulation and biochemical response in South American native species exposed to zinc: Boosted regression trees as novel tool for biomarkers selection. Ecol. Indicators 67, 769–778.

Bradford, M.M., 1976. A rapid and sensitive method for the quantitation of microgram quantities of protein utilizing the principle of protein-dye binding. Anal. Biochem. 72, 248–254.

CAST, 2003. Mycotoxins: risks in plant, animal and human. Potential economic costs of mycotoxins in United States. CAST Task force reports, Council for Agricultural Science and Technology, Ames (IA), pp. 136–142.

Caverzan, A., Casassola, A., Brammer, S.P., 2016. Antioxidant responses of wheat plants under stress. Genet Mol Biol 39, 1–6.

Dantzer, W.R., Pometto, A.L., 3rd, Murphy, P.A., 1996. Fumonisin B1 production by Fusarium proliferatum strain M5991 in a modified Myro liquid medium. Nat Toxins 4, 168–173.

Dastjerdi, R., Karlovsky, P., 2015. Systemic Infection of Maize, Sorghum, Rice, and Beet Seedlings with Fumonisin-Producing and Nonproducing Fusarium verticillioides Strains. Plant Pathol J 31, 334–342.

Desjardins, A.E., Plattner, R.D., 2000. Fumonisin B(1)-nonproducing strains of Fusarium verticillioides cause maize (Zea mays) ear infection and ear rot. J Agric Food Chem 48, 5773–5780.

Devin, S., Burgeot, T., Giamberini, L., Minguez, L., Pain-Devin, S., 2014. The integrated biomarker response revisited: optimization to avoid misuse. Environ Sci Pollut Res Int 21, 2448–2454.

Foyer, C.H., Noctor, G., 2009. Redox regulation in photosynthetic organisms: signaling, acclimation, and practical implications. Antioxid Redox Signal 11, 861–905.

Frisvad, J.C., Larsen, T.O., Thrane, U., Meijer, M., Varga, J., Samson, R.A., Nielsen, K.F., 2011. Fumonisin and ochratoxin production in industrial Aspergillus niger strains. PLoS One 6, e23496.

Glenn, A.E., Zitomer, N.C., Zimeri, A.M., Williams, L.D., Riley, R.T., Proctor, R.H., 2008. Transformation-mediated complementation of a FUM gene cluster deletion in Fusarium verticillioides restores both fumonisin production and pathogenicity on maize seedlings. Mol. Plant-Microbe Interact. 21, 87–97.

Glenz R., Schmalhaus D., Krischke M., Mueller M.J., and Waller F. 2019. Elevated Levels of Phosphorylated Sphingobases Do Not Antagonize Sphingobase- or Fumonisin B1-Induced Plant Cell Death. Plant and Cell Physiology 60, 1109–1119.

Heath, M.C., 2000. Hypersensitive response-related death. Plant Mol. Biol. 44, 321–334.

Heath, R.L., Packer, L., 1968. Photoperoxidation in isolated chloroplasts. I. Kinetics and stoichiometry of fatty acid peroxidation. Arch Biochem Biophys 125, 189–198.

Igarashi D., Bethke G., Xu Y., Tsuda K., Glazebrook J., and Katagiri F. 2013. Pattern-Triggered Immunity Suppresses Programmed Cell Death Triggered by Fumonisin B1. PLOS ONE 8, e60769.

IPCS-WHO, 2000. Fumonisin B1. Environmental Health Criteria 219. In: International Programme on Chemical Safety (Ed.), World Health Organization, Geneva.

Klessig D.F., Choi H.W., and Dempsey D.M.A. 2018. Systemic Acquired Resistance and Salicylic Acid: Past, Present, and Future. Molecular Plant-Microbe Interactions 31, 871–888.

Kwak J.M., Nguyen V., and Schroeder J.I. 2006. The Role of Reactive Oxygen Species in Hormonal Responses. Plant Physiology 141, 323–329.

Loake G., and Grant M. 2007. Salicylic acid in plant defence—the players and protagonists. Current Opinion in Plant Biology 10, 466–472.

Maschietto, V., Lanubile, A., Leonardis, S.D., Marocco, A., Paciolla, C., 2016. Constitutive expression of pathogenesis-related proteins and antioxydant enzyme activities triggers maize resistance towards Fusarium verticillioides. J Plant Physiol 200, 53–61.

Monferran, M.V., Agudo, J.A., Pignata, M.L., Wunderlin, D.A., 2009. Copper-induced response of physiological parameters and antioxidant enzymes in the aquatic macrophyte Potamogeton pusillus. Environ Pollut 157, 2570–2576.

Noctor G., Reichheld J.-P., and Foyer C.H. 2018. ROS-related redox regulation and signaling in plants. Seminars in Cell & Developmental Biology 80, 3–12.

Overmyer K., Brosché M., and Kangasjärvi J. 2003. Reactive oxygen species and hormonal control of cell death. Trends in Plant Science 8, 335–342.

Pan X., Welti R., and Wang X. 2008. Simultaneous quantification of major phytohormones and related compounds in crude plant extracts by liquid chromatography-electrospray tandem mass spectrometry. Phytochemistry 69, 1773–1781.

Presello, D., Iglesias, J., Fernández, M., Fauguel, C., Eyhérabide, G., Lorea, R., 2009. Reacción de cultivares a hongos productores de micotoxinas en maíz. Instituto Nacional de Tecnología Agropecuaria.

Presello, D.A., Iglesias, J., Botta, G., Reid, L.M., Lori, G.A., Eyhérabide, G.H., 2006. Stability of maize resistance to the ear rots caused by Fusarium graminearum and F. verticillioides in Argentinian and Canadian environments. Euphytica 147, 403–407.

Pusztahelyi, T., Holb, I.J., Pocsi, I., 2015. Secondary metabolites in fungus-plant interactions. Front Plant Sci 6, 573.

Rizhsky, L., Hallak-Herr, E., Van Breusegem, F., Rachmilevitch, S., Barr, J.E., Rodermel, S., Inze, D., Mittler, R., 2002. Double antisense plants lacking ascorbate peroxidase and catalase are less sensitive to oxidative stress than single antisense plants lacking ascorbate peroxidase or catalase. Plant J 32, 329–342.

Santiago, R., Cao, A., Butron, A., 2015. Genetic Factors Involved in Fumonisin Accumulation in Maize Kernels and Their Implications in Maize Agronomic Management and Breeding. Toxins (Basel) 7, 3267–3296.

Saucedo-Garcia, M., Gonzalez-Solis, A., Rodriguez-Mejia, P., Olivera-Flores Tde, J., Vazquez-Santana, S., Cahoon, E.B., Gavilanes-Ruiz, M., 2011. Reactive oxygen species as transducers of sphinganine-mediated cell death pathway. Plant Signal. Behav. 6, 1616–1619.

Selin, C., de Kievit, T.R., Belmonte, M.F., Fernando, W.G., 2016. Elucidating the Role of Effectors in Plant-Fungal Interactions: Progress and Challenges. Front Microbiol 7, 600.

Shephard, G.S., Sydenham, E.W., Thiel, P.G., Gelderblom, W.C.A., 1990. Quantitative determination of fumonisins B1 and B2 by high performance liquid chromatography with fluorescence detection. J. Liq. Chromatogr. 13, 2077–2087.

Stone, J.M., Heard, J.E., Asai, T., Ausubel, F.M., 2000. Simulation of fungal-mediated cell death by fumonisin B1 and selection of fumonisin B1-resistant (fbr) Arabidopsis mutants. Plant Cell 12, 1811–1822.

Susca, A., Moretti, A., Stea, G., Villani, A., Haidukowski, M., Logrieco, A., Munkvold, G., 2014. Comparison of species composition and fumonisin production in Aspergillus section Nigri populations in maize kernels from USA and Italy. Int J Food Microbiol 188, 75–82.

Theumer, M.G., Lopez, A.G., Aoki, M.P., Canepa, M.C., Rubinstein, H.R., 2008. Subchronic mycotoxicoses in rats. Histopathological changes and modulation of the sphinganine to sphingosine (Sa/So) ratio imbalance induced by Fusarium verticillioides culture material, due to the coexistence of aflatoxin B1 in the diet. Food Chem Toxicol 46, 967–977.

Wang, X., Wu, Q., Wan, D., Liu, Q., Chen, D., Liu, Z., Martinez-Larranaga, M.R., Martinez, M.A., Anadon, A., Yuan, Z., 2016. Fumonisins: oxidative stress-mediated toxicity and metabolism in vivo and in vitro. Arch. Toxicol. 90, 81–101.

Waskiewicz, A., Bocianowski, J., Perczak, A., Golinski, P., 2015. Occurrence of fungal metabolites--fumonisins at the ng/L level in aqueous environmental samples. Sci Total Environ 524–525, 394-399.

Williams, L.D., Glenn, A.E., Zimeri, A.M., Bacon, C.W., Smith, M.A., Riley, R.T., 2007. Fumonisin disruption of ceramide biosynthesis in maize roots and the effects on plant development and Fusarium verticillioides-induced seedling disease. J Agric Food Chem 55, 2937–2946.

Xing, F., Li, Z., Sun, A., Xing, D., 2013. Reactive oxygen species promote chloroplast dysfunction and salicylic acid accumulation in fumonisin B1-induced cell death. FEBS Lett 587, 2164–2172.

Zhang X., Wu Q., Cui S., Ren J., Qian W., Yang Y., He S., Chu J., Sun X., Yan C., Yu X., and An C. 2015. Hijacking of the jasmonate pathway by the mycotoxin fumonisin B1 (FB1) to initiate programmed cell death in Arabidopsis is modulated by RGLG3 and RGLG4. Journal of experimental botany 66, 2709–2721.

Zhao, Y., Wang, J., Liu, Y., Miao, H., Cai, C., Shao, Z., Guo, R., Sun, B., Jia, C., Zhang, L., Gigolashvili, T., Wang, Q., 2015. Classic myrosinase-dependent degradation of indole glucosinolate attenuates fumonisin B1-induced programmed cell death in Arabidopsis. Plant J 81, 920–933.

Zitomer, N.C., Jones, S., Bacon, C., Glenn, A.E., Baldwin, T., Riley, R.T., 2010. Translocation of sphingoid bases and their 1-phosphates, but not fumonisins, from roots to aerial tissues of maize seedlings watered with fumonisins. J Agric Food Chem 58, 7476–7481.

